# Taking it slow: A positive feedback chloride current in cone photoreceptors supports slow motion detection

**DOI:** 10.64898/2025.12.08.692952

**Authors:** Matthew Yedutenko, Marcus H.C. Howlett, Gabrielle van der Veen, Maarten Kamermans

## Abstract

Photoreceptors gate and shape visual information. Since perception ultimately guides action, the way photoreceptors transform light into neural signals constrains an animal’s behavioural repertoire. Yet, the physiological role of the highly conserved calcium-dependent chloride current in cone photoreceptors has remained elusive. Whereas calcium-activated currents typically provide negative feedback, we identify a counter-intuitive positive feedback mechanism that enhances detection of slow motion. Using genetics, electrophysiology, and behavioural assays in zebrafish, we show that loss of this current exaggerates the biphasic character of cone impulse responses and diminishes their voltage response amplitude to low temporal frequencies (<1 Hz). These changes in photoreceptor properties translate to behaviour, reducing the gain of the optokinetic response to slow-moving stimuli. Mechanistically, the calcium-dependent chloride current acts as a positive feedback loop that sustains the photovoltage response and counteracts strong adaptation within the cone phototransduction cascade. This functional role is conserved across sensory modalities, as demonstrated by similar temporal integration and amplification observed in the odorant system.

## Introduction

Photoreceptors gate and shape visual information. Since perception ultimately guides action, the way photoreceptors transform light into neural signals constrains an animal’s behavioural repertoire. This transformation depends on the characteristics on the involved voltage-gated ion channels that convert changes in the so-called photocurrent (that results from absorption of photons) into changes in the photoreceptor membrane potential. Importantly, the contribution of the various ion channels may differ in different stimulus conditions (1–8). One of the largest, yet most functionally ambiguous, components of this conversion is highly conserved calcium-dependent chloride current (I_Cl[Ca]_) (9–11) that was reported in reptiles (8,12–15), amphibians (9,16,17), fish (18), and mammals (including primates)(19–21).

Typically, the calcium-dependent currents in excitable cells work as a negative feedback regulators, that provide dampening of the response, high-pass filtering of the incoming signals and adaptation(22,23). This is also occurs in the photoreceptors, where calcium-dependent regulation of phototransduction cascade provides adaptation to light intensity(6,24), while feedback from horizontal cells subtracts spatio-temporal redundancies effectively providing high-pass filtering of the input signals(6,25). Thus, one might expect similar functional role of I_Cl[Ca],_ like indeed was sometimes hypothesized(5,15,20). On the other hand depending on chloride reversal potential and similarly to how I_Cl[Ca]_ functions in odorant system(26,27), it was proposed that I_Cl[Ca]_ might also provide positive feedback to amplify input signals (3,5).

However, despite biophysical properties(3,5,9–12,14,16,17,28–36), proteins and genes associated with I_Cl[Ca]_(28,29,37) being well-characterized, investigation of its physiological role in the earlier studies were plagued by non-physiological recording conditions and thus remains unclear. Some studies(8,12,13,16) had way too high (M) chloride concentrations making reversal chloride reversal potential about 100 mV higher than suggested by the measurements (3,5,18,38); others (3,18,19,29,30,35) employed concentrations of calcium buffer (1-5mM EGTA) that were 100 times higher than the endogenous buffer capacity (0.05-0.1 mM EGTA) (35,36) and therefore were effectively blocking that distorted chloride reversal potential, others used supraphysiological calcium buffering that effectively blocked the I_Cl[Ca]_ (35); third (3,29,30,35) extrapolated its role in light signal processing from response to electrical stimuli that were driving photoreceptor membrane with potentials 20-60 times higher than the light responses(2,39–41). The work by Frederiksen et al. (21), actually, describes function of I_Cl[Ca]_ in physiological conditions in rods, however, as a light stimuli they employed short flashes of light, instead of dynamic stimuli characteristic for natural scenes(42).

Here, we provide the definitive mechanistic resolution of this long-standing functional mystery. We generated a zebrafish strain with a general knock-out of the Anoctamine 2 (Ano2) gene that encodes the I_Cl[Ca]_ -conducting ionic channel TMEM16B (Figure 1 B,F). We first established the behavioural consequence of the knock-out, finding that mutant (Ano2^−/−^) zebrafish has decreased amplitude of the optokinetic response (the ability to visually track moving stimuli, OKR), especially at low velocities. This behavioural link suggests that I_Cl[Ca]_-mediated signals are crucial for animal’s motion sensitivity. To explain this phenotype, we recorded voltage light responses of cone photoreceptors with physiological E_Cl_ and concentration of the calcium buffer.

**Figure 1.**
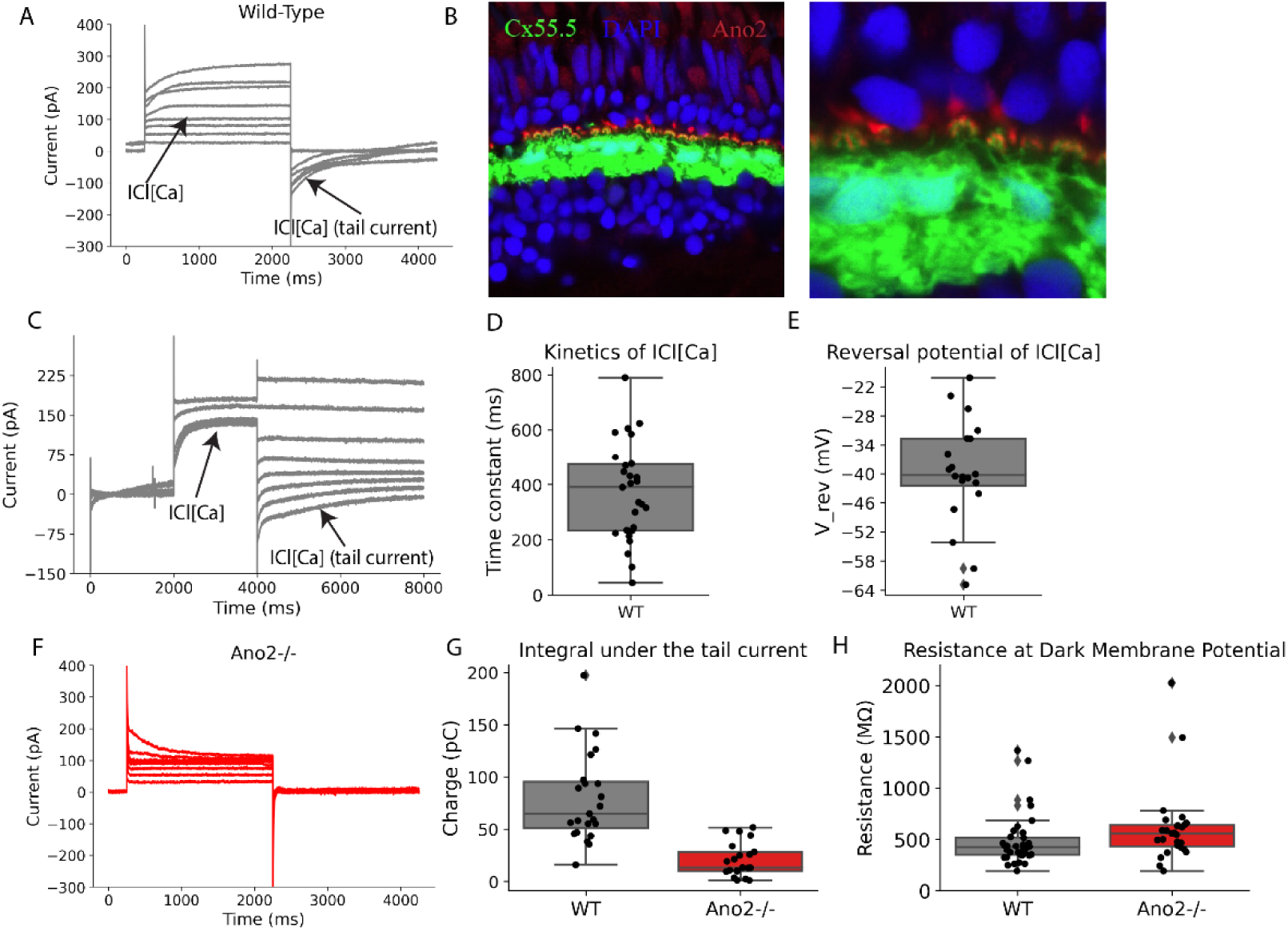
Basic properties of I_Cl[Ca]._ A. An example of the currents recorded in a response to change in the membrane potential in WT zebrafish cones. The cells were depolarized from a holding potential of −70 mV to various potentials (steps of 10 mV) for 2000 ms and then stepped back to −70 mV. Left arrow – contribution of the I_Cl[Ca]_ to the current recorded upon depolarization. Right arrow – tail current. B. Fluorescent microscopy images of the Outer Plexiform Layer (left) and the magnified image of the cone synaptic terminals (right). Green – GFP-labelled Connexin 55.5 in horizontal cells, blue – DAPI that stains cell nuclei, red – antibody to Ano 2. C. An example of tail currents (right arrow) recorded in WT zebrafish cones after stepping to various holding potentials (steps of 10mV) from a holding potential of 10mV (left arrow). D. Boxplot of the estimated time constant of I_Cl[Ca]_ tail currents in the WT zebrafish cones. E. Boxplot of the I_Cl[Ca]_ reversal potentials in WT zebrafish cones determined using the protocol shown in panel C. F. An example of the protocol shown in panel A repeated in an Ano2^−/−^ zebrafish cone. G. Boxplots comparing the integral under the tail current in WT (grey) and mutant (red) zebrafish cones. H. Boxplots comparing the input resistance of WT (grey) and mutant (red) zebrafish cones at their resting membrane potentials in the dark.

Contrary to the expected negative-feedback role, our data reveal that under physiological conditions I_Cl[Ca]_ in cone photoreceptors acts as a novel positive-feedback low-pass filter. Loss of I_Cl[Ca]_ enhanced the biphasic nature of the cone impulse response and reduced the response gain at temporal frequencies below 1.3 Hz (Figure 4). Effectively, the activation of I_Cl[Ca]_ compensates for negative feedback inherent in the adaptation within the phototransduction cascade. By enhancing responses to these frequencies, the I_Cl[Ca]_-mediated feedback loop restores their Signal-to-Noise Ratio (SNR), retaining information about slowly changing features of the visual scene, that is relevant for, for example, detection of slow motion. Notably, the effect of I_Cl[Ca]_ can be mitigated by decrease in stimulus contrast.

The positive feedback and low noise signal amplification by I_Cl[Ca]_ observed here in cone photoreceptors is consistent with the function reported in the olfactory system(26,27,43). Therefore our study provides necessary evidence for a principle of low noise signal amplification by I_Cl[Ca]_ that is conserved across the primary receptors in different sensory modalities. Critically, we demonstrate the functional necessity of this mechanism by linking Ano2^−/−^ knockout to tangible consequences for visual-guided behaviour.

## Results

### Basic properties of ICl[Ca]

Figure 1A shows recordings of I_Cl[Ca]_ in WT cones (Figure 1B). The cone membrane potential was stepped for 2000 ms to various potentials from a holding potential of −70 mV. Depolarization induces a slowly activating outward current (left arrow) and a large tail current when the potential was stepped back to the holding potential (right arrow). To prevent interference of other voltage-gated currents, we used the tail current to estimate two basic properties of I_Cl[Ca]_: its time constant and reversal potential.

To estimate the time constant of I_Cl[Ca],_ we fitted tail currents with a single exponential function and found a time constant of 373 ± 34 ms (n=27, Figure 1D). In the frequency domain, this is equivalent to a first-order low-pass filter with a 3db cut-off frequency of ∼0.4Hz, suggesting that the activation/deactivation of the channel mostly affect responses to temporal frequencies up to 1 Hz.

To determine the reversal potential of I_Cl[Ca]_, we recorded the tail current at different holding membrane potentials and interpolated the value at which the tail current became zero. The reversal potential found was −39 ± 2.5 mV (n=19, Figure 1E), which is slightly more depolarized than the average resting membrane potential of WT cones in the dark (−45 ± 0.9 mV, n=17).

### TMEM16B is a molecular correlate of Icl[ca] in zebrafish cone photoreceptors

To determine the effectiveness of our KO strategy, we repeated the protocol from Figure 1A on mutant cones (Ano2^−/−^, Figure 1F) and compared their recordings to those of WT cones. Qualitatively, the mutant cones exhibited markedly reduced slow outward and tail currents.

To quantify the effect of the mutation, we compared the integral under the tail current after depolarization to 10mV in WT and mutant zebrafish cones (Figure 1G). In mutant cones, the integrated tail current was 4 times smaller (WT:80 ± 9.3 pC; Mutant: 20 ± 4pC, n=21; p=5e-7, N_DOF_=41), indicating that I_Cl[Ca]_ was absent.

If the TMEM16B channels are absent in mutant and present in WT, then one would expect that the input resistance of the cones at their dark resting membrane potential will be higher in the mutant compared to WT. Indeed, WT cones had a significantly lower membrane resistance than Ano2^−/−^ cones: 482 ± 42 MΩ vs. 608 ± 71MΩ, N_DOF_=58, p=0.006 (Figure 1H).

### ICl[Ca] in cones allows zebrafish to better track slowly moving stimuli

To test whether an absence of the I_Cl[Ca]_ in mutant fish has any consequences for their visually-guided behaviour, we recorded the optokinetic response of immobilized zebrafish larvae (7 d.p.f.) to moving gratings (44), (Figure 2A, B). We tested 3 spatial frequency, and 6 different angular velocities.

**Figure 2.**
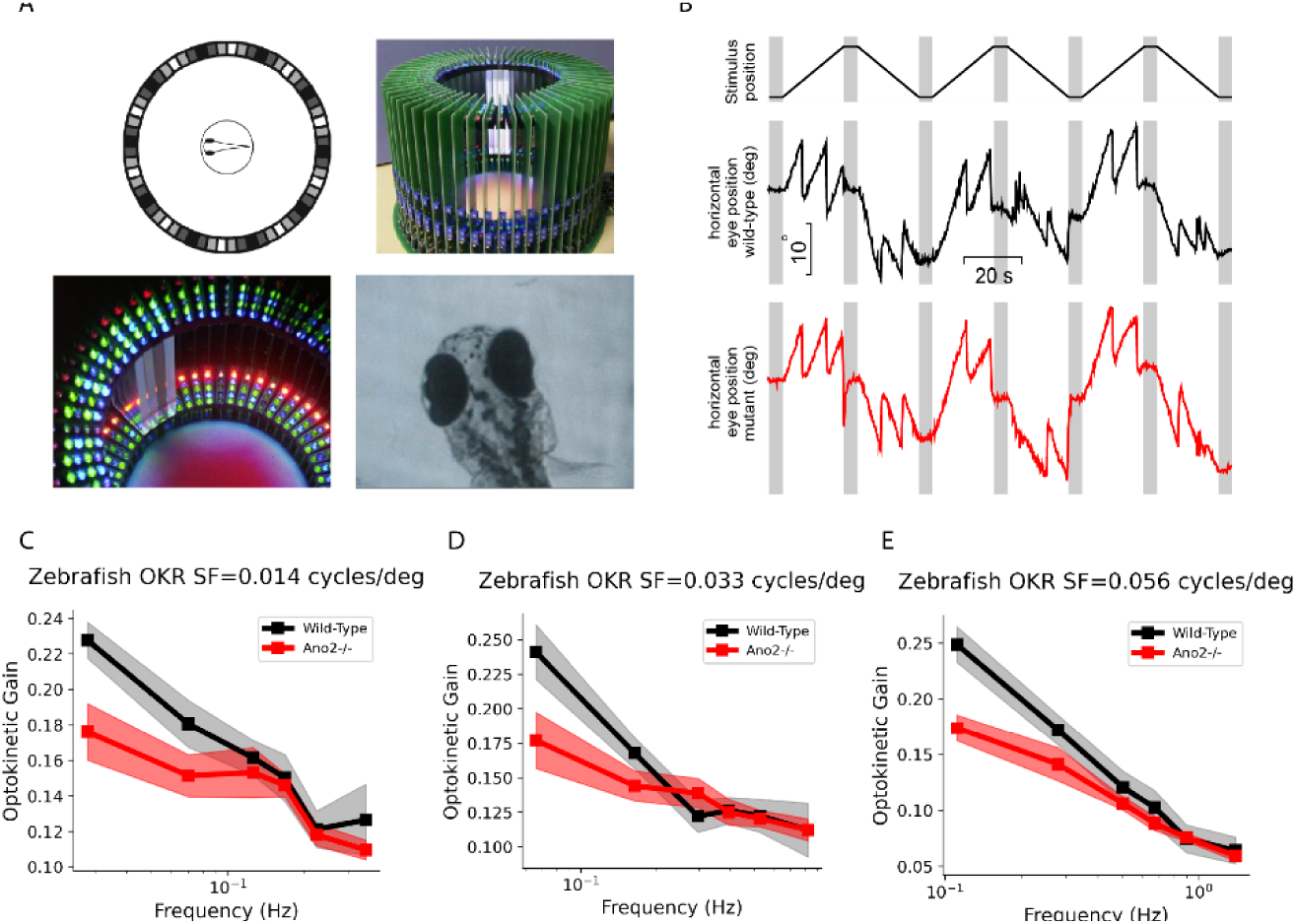
Optokinetic response in WT (black) and Ano2^−/−^ (red) zebrafish larvae. A. Experimental setup (from left-to-right): diagram of the setup, recording chamber, LED lights that create moving pattern, frame from an eye-tracking video.. B. An example of the measured angular velocity of OKR eye movements. C-E Optokinetic gain plotted for three spatial frequencies across six temporal frequencies. Line – mean value, shading – standard error of the mean.

In Figures 2C-E we plotted the optokinetic response gain, as a function of stimulus temporal frequency, for each of the spatial frequencies. All plots show that the mutant zebrafish exhibited reduced optokinetic gain at temporal frequencies between 0.02 – 0.2 Hz, consistent with the dynamics of I_Cl[Ca]_ (Figure 1). To quantify the genotype’s effect, we applied a linear mixed-effects model to data collected at a stimulus velocity of 2°/s, with genotype and spatial frequency as fixed effects and individual fish as a random effect. The model revealed that, on average, WT animals have a 36% higher optokinetic gain compared to mutant animals (Wald Z test(45), p=0.005, N_observations_=43, N_groups_=17). This implies that activation of I_Cl[Ca]_ enhances the ability of zebrafish to track slowly moving stimuli. To summarize, Figure 2 indicates the importance of I_Cl[Ca]_ for zebrafish visually-guided behaviour.

### Absence of ICl[Ca] affects cone light responses

Next, we examined whether the altered OKR in mutants (Figure 2) could be attributed to differences in the light response characteristics of zebrafish. To this end, we recorded voltage responses of Ano2^−/−^ and WT cones (Figures 3B&C) to light decrements (negative contrast) and increments (positive contrast) with magnitude of 0.8 Weber units (change in light intensity relative to baseline light intensity, see Methods for details). The stimulus waveform is shown on Figure 3A (note the sign reversal of the light response). Although the peak amplitude of the responses of WT and Ano2^−/−^ cones did not differ significantly (negative contrast: WT +2.4 ± 0.2 mV, mutant 2.4 ± 0.3 mV, p=0.98; positive contrast: WT −0.9 ± 0.08 mV, mutant –1.1 ± 0.1 mV, p=0.2, N_DOF_=20), their responses kinetics were distinct.

**Figure 3.**
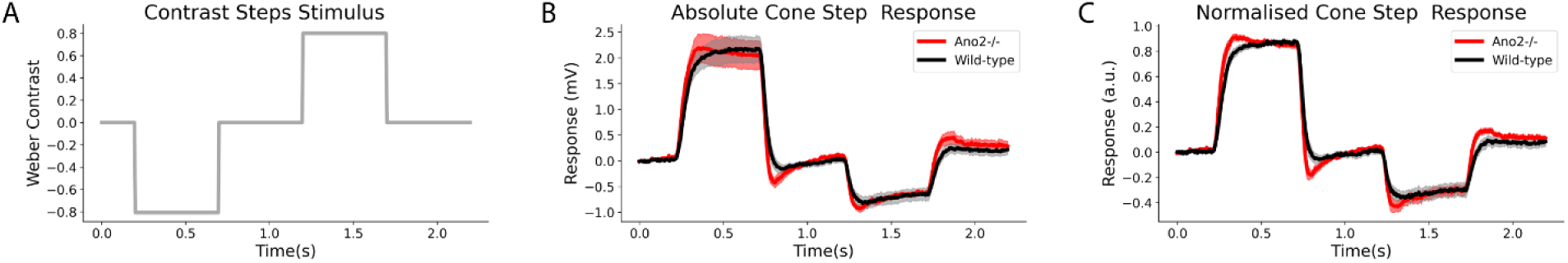
Zebrafish WT (black) and Ano2^−/−^ (red) cone photoreceptor responses to steps of positive and negative contrast. Solid lines – mean values, shadings - ±SEM. A. – Stimulus as function of time. B. Absolute voltage response of WT and Ano2^−/−^ cones. C. Normalized voltage responses of WT and Ano2^−/−^cones.

To visualize this difference in kinetics better, we normalized the step responses to their peak response amplitude (Figure 2C). On average, responses of WT zebrafish cones to negative contrast steps peaked ∼375 ± 25 ms after the stimulus onset (N=13), closely matching the kinetics of the I_Cl[Ca]_ tail current kinetics shown in Figure 1. Mutant cone responses peaked significantly earlier, at ∼ 145 ± 35 ms (N=9), a ∼230ms difference (N_DOF_=20, p= 2e-4). For the positive contrast steps, WT cone responses peaked ∼100ms later than it did for the mutants (WT: 200 ± 32ms, Ano2^−/−^ : 100 ± 9ms, p=0.017, N_DOF_=20).

Ano2^−/−^ cones also showed a greater degree of response rollback, measured as the ratio between the amplitude of a cone’s voltage response at the end of the contrast step and its peak response. For negative contrast steps, the rollback of Ano2^−/−^ cones was almost 2 times greater than for WT cones: 21 ± 2.7% vs. 11 ± 2%, p=0.008, N_DOF_=20. For voltage responses to positive contrast step, the rollback of Ano2^−/−^ cones was ∼1.5 times greater than for WT cones: 36 ± 3.6% vs. 23 ± 3%, p=0.013, N_DOF_=20.

To summarize, in cone photoreceptors, I_Cl[Ca]_ activation delays the peak voltage response to steps of both positive and negative contrast and reduces the magnitude of response rollback.

### Activation of ICl[Ca] makes cone voltage light responses less biphasic

To further characterize these differences in cone kinetics, we recorded Ano2^−/−^ and WT cone responses to a sum of sinewaves light stimulus (Figure 4A). This is a classical system identification technique that tests a system’s response to multiple frequencies at the same time (46). To mimic the power spectrum of naturalistic scenes, we selected sinewave frequencies such that most of the stimulus power was concentrated in the lower frequency end of the spectrum (see methods for the details). We focused this part of our analysis on short-wavelength sensitive cones (S-cones), as they have a higher passive membrane resistance than mid-(M) and long-(L) wavelength sensitive cones (453 ± 29 MOhm for S-cones vs 353 ± 27 MOhm for M-cones, p=0.0245, N_DOF_=16, unpublished observation). This higher resistance amplifies the effect of I_Cl[Ca]_ on the S-cone membrane potential, thereby emphasizing difference between Ano2^−/−^ and WT cones.

**Figure 4.**
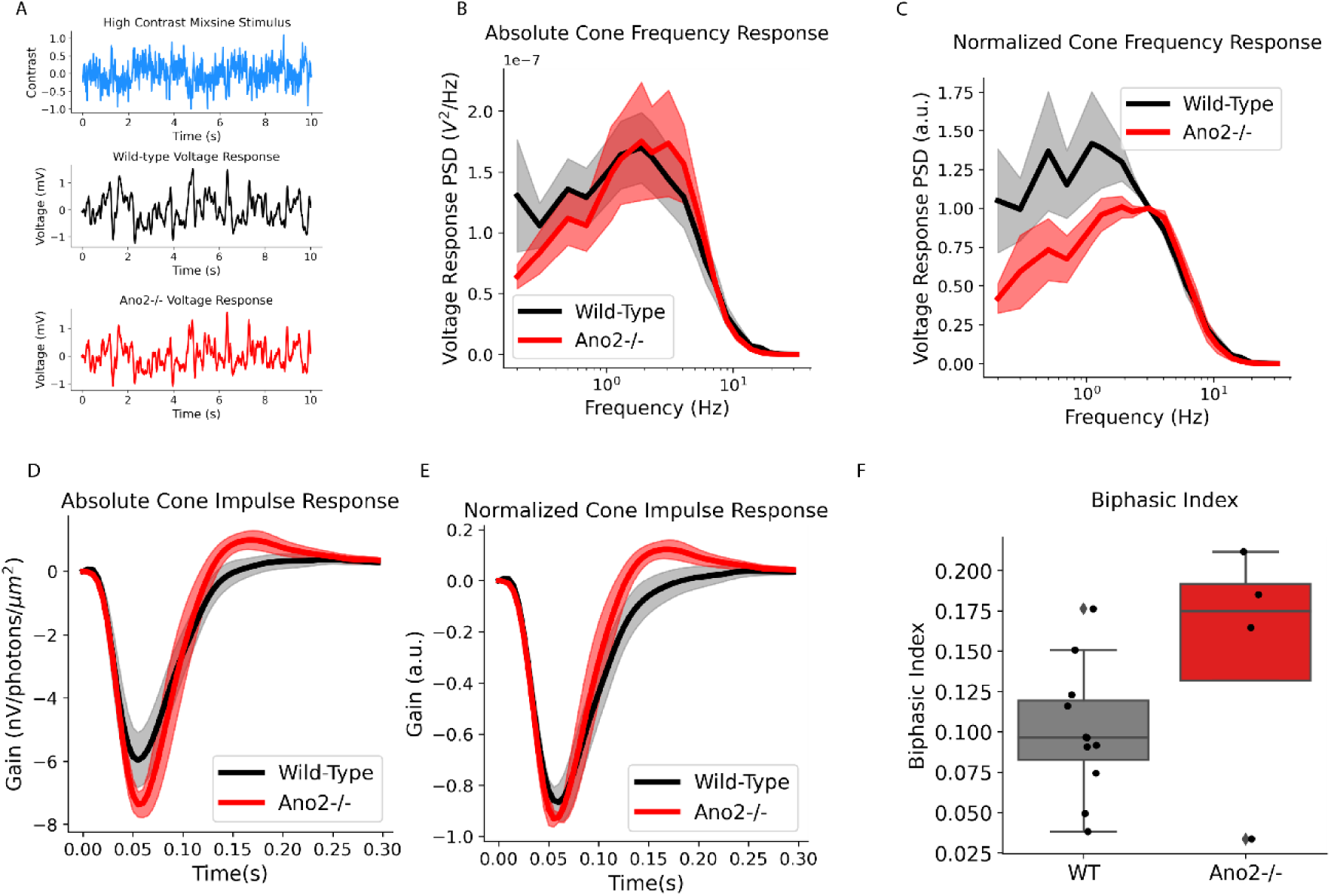
WT (black) and Ano2^−/−^(red) zebrafish S-cone signal transfer properties. Solid lines – mean values, shadings -±SEM A. Top. Sum of Sinewaves Stimuli. Middle. An example trace of WT cone voltage response. Bottom. Example of Ano2^−/−^ cone voltage response. B. Absolute Frequency response of WT and Ano2^−/−^ S-cones. C. Normalized Frequency response of WT and Ano2^−/−^ S-cones. D. Absolute Impulse Response of WT and Ano2^−/−^ S-cones. E. Absolute Impulse Response of WT and Ano2^−/−^ S-cones. F. Boxplot of the biphasic index for WT (grey) and Ano2^−/−^ S-cones (red).

Using fast Fourier transform, the responses of cones were plotted as a function of stimulus frequency (Figure 4B&C). The curves for both WT and Ano2^−/−^ show bandpass characteristics but still differ in their frequency dependence. Figures 4B&C showed WT cones had larger absolute (Figure 4B) and relative (Figure 4C) response amplitude at frequencies below 1Hz, consistent with the kinetics of I_Cl[Ca]_ (Figure 1) and frequency range over which WT zebrafish showed higher OKR gain than Ano2^−/−^ zebrafish (Figure 2).

To more parsimoniously characterize differences in temporal signal processing between WT and mutant cones, we computed their impulse response functions (Figures 4D&E) by making an inverse Fourier transform of the frequency response curves shown in Figure 4B. Both absolute (Figure 4D) and normalized (Figure 4E) impulse responses revealed no statistically significant difference in peak amplitude or in the integration time of the primary lobe, which reflects the initial response to stimulus onset. However, mutant cones exhibited markedly more biphasic impulse responses, with nearly a two-fold increase in the biphasic index (0.19 ± 0.014 in mutant vs. 0.10 ± 0.013 in wild type; NDOF = 11, p = 0.001; Figure 4F). The biphasic index, calculated as the ratio of the second to the first peak amplitude, increases when responses become more transient and less sensitive to slow stimulus changes.

In summary, comparisons of the impulse responses between mutant and WT cones (Figure 4) shows that I_Cl[Ca]_ activation reduces the secondary lobe of the impulse response, resulting in a more low-pass frequency response.

### ICl[Ca] activation increases the fraction of information encoded by cone photoreceptors at low temporal frequencies

From Figures 4 B&C we learned that activation of I_Cl[Ca]_ increases the amplitude of photoreceptor light responses to temporal frequencies below 1.3 Hz. But is this a simple passive amplification that equally affects both the signal and noise components of a cone’s response, or is it an active process that selectively enhances the signal component? A useful metric to address this question is the information transfer rate, which is proportional to the ratio between signal and noise power (i.e. Signal-to-Noise Ratio or SNR; see Methods for the mathematical details).

In Figures 5A and 5B we plotted the cone information transfer rate as the function of the stimulus temporal frequency. The normalized spectra (Figure 4B) indicate that compared to WT cones, mutant cones (red trace) allocated a lesser fraction of their information channel capacity towards frequencies lower than 1.3Hz. This is consistent with the more biphasic impulse response of mutant cones. (Figure 4). Quantitatively, activation of I_Cl[Ca]_ amplified the relative information transfer of WT cones for frequencies below 1.3 Hz by 30% compared to mutant cones (Figure 4C, p=0.046, N_DOF_ =12). This frequency range is consistent with the 373 ms time constant of I_Cl[Ca]_ activation reported in Figure 1. To summarize, activation of I_Cl[Ca]_ in zebrafish S-cones enhances information transmission at frequencies below 1.3Hz.

**Figure 5.**
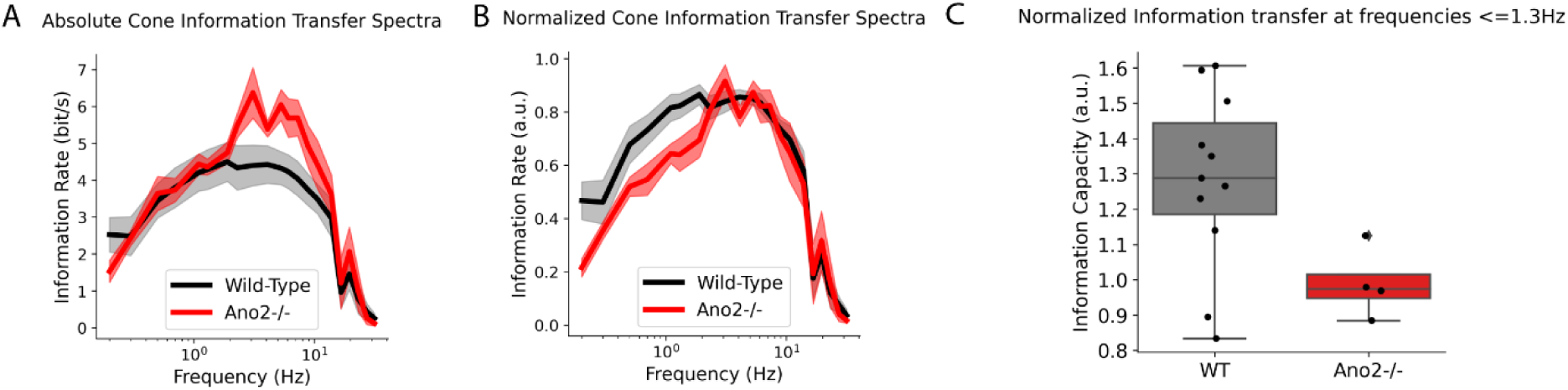
Information transmission in WT (black) and Ano2^−/−^ (red)^−^ zebrafish S-cones. Solid lines – mean values, shadings - ±SEM A. Absolute Cone Information Transfer Spectra. B. Normalized Information Transfer Spectra. C. Normalized Information Channel Capacity at frequencies below 1.3Hz.

### ICl[Ca] Modulation of Cone Signals Is Diminished at Low Contrast

Next, we tested whether the effects of I_Cl[Ca]_ on cone signals and information transfer properties depends on the stimulus contrast. To do so, we recorded cone voltage responses to a low contrast sum of sinewaves light stimulus (Figure 6A). The variance of light intensity around the mean in this low-contrast stimulus was 5 times lower than that of the high-contrast stimulus used previously. From these recording, we derived the frequency response curves (Figure 6B-6C), the impulse response functions (Figure 6D-6E) and the information spectra analysis (Figure 6G-6H).

**Figure 6.**
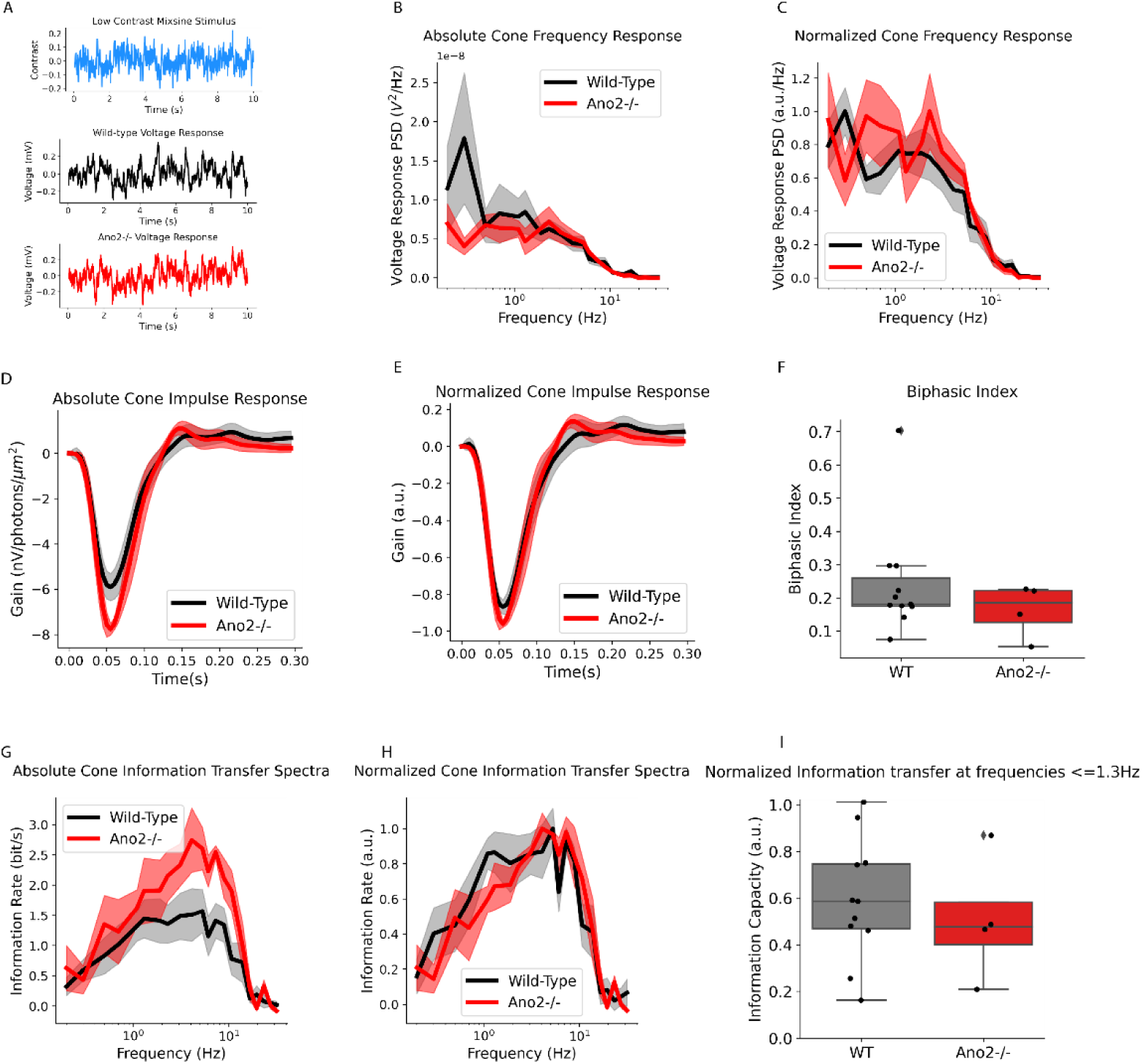
Signal and information transfer properties of WT (black) and Ano2^−/−^ (red) zebrafish S-cones for the low contrast light stimuli. Solid lines – mean value, shading - ±SEM. A. Top. Low contrast sum of sinewaves stimulus. Middle. An example of WT zebrafish S-cone voltage response. Bottom. An example of Ano2^−/−^ S-cone voltage response to low contrast light stimulus. B. Absolute S-cone frequency responses. C. Normalized frequency responses. D. Absolute S-cone impulse response. E. Normalized S-cone impulse response. F. Biphasic Index for low contrast G. Absolute S-cone information transfer spectra. H – Normalized cone information transfer spectra. I – Normalized information transfer at frequencies < 1.3Hz.

Figures 6B-C show that I_Cl[Ca]_ has little effect on frequency response curves at low contrasts. The apparent bump in the WT curve in Figure 6B is due to a single cell that exhibited an outlier response at that specific frequency, thereby elevating the mean. Similarly, Figures 6D-F show that, under low-contrast conditions, the relative amplitude of the secondary lobe of the impulse response does not differ between mutant (red trace) and WT (black trace) cones (0.20 for WT cones, 0.16 for mutant cones, N_DOF_=11, p=0.96). Consistent with these finding, the information transfer rates in mutant and WT cones were similar under low contrast conditions (Figure 6G-I, cf. Figure 5), particularly at frequencies below 1.3Hz (Figure 6I; p=0.538, N_DOF_=12). Also note qualitative similarities between low contrast information transfer plots of both animals and high mutant information transfer plots (Figure 5).

To summarize, under low contrast stimulus conditions, I_Cl[Ca]_ had little effect on the cone’s signal (Figure 6 B-F) and information transfer properties (Figure 6 D-I).

## Discussion

In this study, we systematically investigated the role of I_Cl[Ca]_ in cone signal processing and its contribution to visually guided behaviour in animals. We generated an Ano 2 knock-out zebrafish and showed that these zebrafish lack I_Cl[Ca]_ (Figure 1) (28,31,34,37,47). We compared visually guided behaviour (OKR) in WT and Ano2^−/−^ zebrafish and found that knocking out I_Cl[Ca]_ impaired their ability to track slowly moving stimuli (Figure 2).

Next, using physiological concentrations of calcium buffer (35,36), Ca^2+^ concentrations corresponding to endogenous levels found in dark-adapted cone photoreceptors (48), and an intrapipette Cl^−^ concentration similar to that estimated for goldfish cones (18), we compared voltage responses of mutant and WT zebrafish cones. We found that activation of I_Cl[Ca]_ makes their impulse responses less biphasic (Figure 4 D-E) and causes cones to behave more like integrators. As a result, activation of I_Cl[Ca]_ increases the cone response gain to frequencies below 1.3Hz (Figure 4 B-C). These changes in response behaviour are also reflected in the cone information coding properties (Figure 5), with WT cones dedicating more relative information channel capacity to temporal frequencies below 1.3Hz than mutant cones. These differences were no longer visible when we lowered the contrast fivefold (Figure 6).

Together, our data suggest that I_Cl[Ca]_ plays role in visually guided behaviour in zebrafish by shaping signal transfer properties of cone photoreceptors.

While these findings underscore the role of cone I_Cl[Ca]_ in shaping photoreceptor signal processing, additional mechanisms may also contribute to the observed deficits in visually guided behaviour. Since Ano2 was knocked out globally, rather than selectively in cone photoreceptors, OKR differences between mutant and WT fish may reflect effects beyond the absence of I_Cl[Ca]_ in cones. For instance, I_Cl[Ca]_ is also expressed in other retinal cell types (32,49) and in the inferior olive, which is involved in motor learning and control(50). However, photoreceptors represent the initial site of visual signal processing, and loss of information at this stage cannot be restored by downstream circuits. Thus, regardless of downstream processing, OKR performance remains fundamentally constrained by the fidelity of photoreceptor output. The similarity between photoreceptor frequency responses (Figure 4) and OKR performance (Figure 2) illustrates this constraint.

The described here function of I_Cl[Ca]_ as a low noise positive feedback integrator is conserved within and across different sensory modalities. For instance, in rod photoreceptors presence of I_Cl[Ca]_ decreases dark noise in rods and rod bipolar cells. (21). While in the odorant system, I_Cl[Ca]_ amplifies response to odorant stimulus by corresponding primary neurons (26,43), extends rise time of their response (26,27) and decreases time required by an animal to find a source of smell(26,43) almost two-fold. Taking all this together, these studies reveal I_Cl[Ca]_ as a conservative mechanism for improving signal-to-noise ratio of low temporal frequency signals in primary neurons across different sensory modalities.

### Electrophysiological properties of Icl[ca]

Analysis of the electrophysiological properties of mutant and WT cones revealed two interesting findings: i) the resistance of the dark resting membrane-potential and ii) the reversal potential of I_Cl[Ca]._

Figure 1H shows that mutant cones exhibit ∼26% higher membrane resistance compared to WT cones. This difference is expected due to the absence of an ion channel. However, what is truly intriguing is that, despite the higher membrane resistance—which typically implies a longer time constant—there was no significant difference in the voltage light response integration time between WT and mutant S-cones (WT: 54.8 ± 3.7 ms, Mutant: 55.5 ± 5.5 ms ,p=0.85, N_DOF_=13). This apparent paradox arises because, although presence of TMEM16B reduces the photoreceptor’s membrane resistance, the slow activation of I_Cl[Ca]._ extends the time over which the input signal is integrated, effectively offsetting the expected reduction in integration time.

Along the same lines, the higher dark membrane resistance in mutants would be expected to increase the amplitude of their responses. However, no significant differences were observed between WT and mutant cones in the peak amplitudes of either step responses or impulse responses. We propose that while I_Cl[Ca]_ initially reduces the cone’s response amplitude, its delayed activation subsequently enhances the overall response. Thus, for instance, although responses of WT cones to steps of negative contrast peak later (with a time-to-peak being similar to the time constant of I_Cl[Ca]_), the maximal amplitude does not change. If this interpretation is correct, then a difference in gain between WT and mutant cones should emerge at low contrast, where the smaller voltage fluctuations barely activate I_Cl[Ca]_ (Figure 6). Indeed, under these conditions, mutant cones showed an average impulse response gain ∼17% higher than that of WT cones, although this difference was not statistically significant (p = 0.3), possibly due to the high variability of noisier responses at low contrast and the limited sample size.

Interestingly, the recordings rod photoreceptors(21) showed that Ano2^−/−^ rods indeed had considerably higher amplitude of light responses. However, in the ref. (21) authors employed different stimulus protocol: 10ms light flashes instead of 400ms contrast flashes (Figure 3) or sum of sinewaves stimuli employed here (Figure 4). The difference in stimuli with, presumably, higher conductance of TMEM16B in rod photoreceptors can explain difference between our results and results obtained in ref. (21). On the other hand, electro-olphactogram recordings in Ano2^−/−^ mice(43) shows that prolonged activation of I_Cl[Ca]_ leads to increased response amplitude in the neurons of odorant system, consistent with our interpretation of the role of I_Cl[Ca]_ in cone photoreceptors light response.

For the reversal potential of I_Cl[Ca],_ Figure 1E shows that, on average, the I_Cl[Ca]_ tail currents reversed at around –39mV. This is slightly more depolarized than the dark resting membrane potential (−45mV) and more depolarized than the calculated chloride reversal potential (E_Cl_ = - 55mV). The former suggests that activation of I_Cl[Ca]_ depolarizes the cone membrane potential, consistent with findings obtained using perforated patch clamp techniques that preserve the endogenous intracellular environment(3,21). The latter is particularly interesting because it was previously assumed that E_Cl_ largely determined the reversal potential of I_Cl[Ca]_, with minimal contributions from other anions(9). However, our results suggest that, under physiological calcium and buffering conditions in zebrafish cones, the TMEM16B channel is not exclusively permeable to chloride ions. The more depolarized reversal potential indicates that other charge carriers may also contribute to the current, as previous studies on TMEM16B channel permeability have shown (reviewed in ref. (37)).

### ICl[Ca] modulation as a positive feedback loop

Our results show that, under physiological conditions, activation of I_Cl[Ca]_ acts as a positive feedback loop that enhances responses to low (∼<1.3 Hz) temporal frequencies in the zebrafish visual system (Figures 2-6).

On a mechanistic level, the influence of I_Cl[Ca]_ on zebrafish vision is realized through the following process. A change in the calcium current (I_Ca_) activates the I_Cl[Ca]_ channel, which in turn adds a slow, low-pass filtered version of that signal back into the photovoltage. When measuring the cone photovoltage step response, this feedback causes a delayed peak in the WT cone’s step response (∼375ms, compared to ∼145ms in mutants) and a reduced amplitude of the response rollback (Figure 3). This reduced rollback response is paralleled by a significantly lower biphasic index in WT cones compared to mutants (nearly half as large), which indicates reduced suppression of slowly varying stimuli (Figure 4).

In the frequency domain, this addition of a low-passed signal is observed as an increase in response amplitude at low (∼<1.3 Hz) temporal frequencies (Figure 4). Consequently, since optokinetic response (OKR) temporal frequencies are all < 1Hz, WT cones provide downstream circuitry with a higher output signal amplitude than mutant cones. As a result, WT zebrafish can better track slowly moving stimuli (Figure 2).

Although inherently unstable, positive feedback loops are widely used in biological systems for signal amplification (51). To make positive feedback useful one needs to stabilize it. Namely, it is necessary to bound the output to prevent runaway effects, and to suppress excessive noise amplification by the feedback loop.

For I_Cl[Ca]_, stabilization of the positive feedback loop arises in part because its amplitude is constrained by the I_Cl[Ca]_ reversal potential during depolarization, and by calcium channel conductance during hyperpolarization. For the depolarizing inputs, I_Cl[Ca]_ drives the cone membrane potential towards the reversal potential. However, the closer the cone potential is to the I_Cl[Ca]_ reversal potential, the smaller the current’s driving force becomes. Hence, as the current approaches its reversal potential its effect on the membrane potential naturally decreases. Moreover, if a large decrease in light intensity depolarizes the cone membrane potential such that it crosses the I_Cl[Ca]_ reversal potential, the current will become hyperpolarizing, and thus start to work as a negative feedback loop. For hyperpolarizing inputs, the resulting drop in membrane potential closes calcium channels, reducing Ca^2+^ influx and thereby limiting I_Cl[Ca]._

With respect to noise mitigation, Figures 5 and 6 suggest that the I_Cl[Ca]_ positive feedback loop does not adversely affect cone signal processing. When light stimulus contrast is high, WT cones encode a proportionally greater share of their signal at lower temporal frequencies compared to mutant cones. This suggests that the I_Cl[Ca]_ feedback loop amplifies the signal more effectively than it amplifies noise.

We propose that in cone photoreceptors, noise mitigation arises from two factors: the slow kinetics of ICl[Ca], which provide a form of temporal averaging to smooth out noise (Figure 1), and the thresholding of input signals. Evidence for signal thresholding comes from cone responses to low-contrast stimuli (Figure 6). Under these conditions, WT and mutant cones transmit similar relative proportions of information across their signalling frequency range, suggesting that the ICl[Ca] feedback loop is largely inactive when input signals are weak. However, as stimulus contrast increases (leading to higher SNR (51,52)), the feedback loop becomes more engaged and exerts a greater influence on cone signal processing, actively shaping the cone’s voltage response (Figures 4,5)

### How is ICl[Ca] integrated in the cone visual pathway?

By acting as a positive feedback loop, the I_Cl[Ca]_ plays a role in retaining information about slowly changing stimulus components. Its importance is obvious for processes requiring information about slow signal features – such as global motion for the OKR response. However, global motion is only one specific type of computation. What are the more general effects of the I_Cl[Ca]_ activation?

The paramount feature of cone photoreceptors is their ability to adapt to changes in light intensities that span 7 decades of magnitude. As a result of this adaptation, the amplitude of photoreceptor responses to temporal contrast is almost independent of the ambient light intensity. This adaptation occurs within the phototransduction cascade and is modulated through a series of Ca-dependent negative feedback loops, each with its dynamics ranging from ∼10ms to ∼10 seconds (53–55). Together, these feedback loops attenuate cone responses to frequencies below 1 Hz. This attenuation is counteracted by the activation of I_Cl[Ca]_, which restores low-frequency components in the cone output.

Why do the lower frequencies need to be retained? Even with adaptation mechanisms in the phototransduction cascade, increases in ambient light intensity tend to hyperpolarize the cone membrane potential(7), and reduce intracellular calcium levels(48). Photoreceptors signal changes in light intensity by modulating vesicle release rates relative to a baseline tonic release. However, reduced membrane potential and calcium influx decrease both the tonic release rate and the pool of available synaptic vesicles(56,57). Because synaptic release is inherently stochastic, reduced vesicle availability and a lower mean release rate amplify the relative impact of random release events, ultimately degrading the SNR of the synaptic output (57). Under these conditions, improving the output’s SNR requires integrating signals over longer time intervals. This is where the slow signal amplification mediated by I_Сl[Сa]_ becomes essential—it preferentially enhances responses to low temporal frequencies, thereby improving SNR.

## Methods

### Animal care

This study was conducted in compliance with the guidelines and approval of the Competent Authority (CCD-license AVD801002017830), the ethical committee of the Royal Netherlands Academy of Arts and Sciences (DEC-KNAW), and the Animal Welfare Body of the Netherlands Institute for Neuroscience (IvD-NIN), following the European Communities Council Directive 2010/63/EU. Wild-type zebrafish (*Danio rerio*, TL strain) were sourced from the Zebrafish International Resource Centre (Eugene, OR, USA; NIH-NCRR grant #RR12546). Both male and female fish were housed in aquaria maintained at a temperature of 28–28.5°C under a 14-hour light and 10-hour dark cycle. They were fed two to three times daily with a combination of live *Artemia nauplii* (INVE Technologies, Dendermond, Belgium; INVE Aquaculture, Salt Lake City, UT, USA) and dry food (Luckstar feed #0.2; Catvis, Netherlands). Zebrafish breeding occurred naturally, triggered by the onset of light. Embryos were reared at 28–28.5°C in E3 medium containing 1 × 10⁻⁵ % methylene blue (Sigma, St. Louis, MO, USA) under standard conditions. The study utilized wild-type zebrafish and mutant line Ano^2-/-^.

### Mutant generation

The CRISPR/Cas9 genome editing technique was used to generate knockout zebrafish (58). Guide RNAs targeting the Ano2 locus were designed using the online tool CHOPCHOP. The target site was located in exon 16 (Figure 7) and had the following target sequence:

GGATGAAATCTATGGCA**\***TGG**TGG** (**\***=cut, **TGG**=pam sequence)

**Figure 7.**
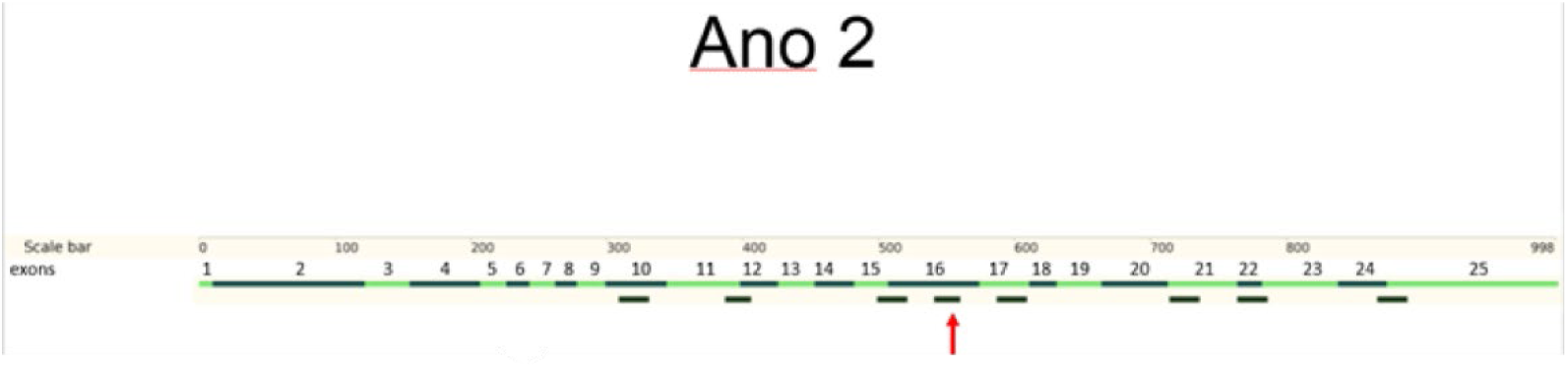
Injection site for the stop codone.

Guide RNAs targeting the Ano2 loci were designed using an online webtool named CHOPCHOP.

Stop codon cassette consisted of an oligonucleotide sequence designed to introduce stop codons in all 3 reading frames (GTCATGGCGTTTAAACCTTAATTAAGCTGTTGTAG) flanked by 20 nucleotide homologue sequences.

Guide RNA (gRNA) and Cas9 mRNA for zebrafish injections were synthesized following the approach described by Gagnon et al. (2014). Briefly, after annealing the oligonucleotides, the resulting DNA templates were used to synthesize gRNA using the Ambion MEGAscript T7 Transcription Kit. Cas9 mRNA was synthesized from NotI-linearized pCS2-nCas9n plasmid DNA (Addgene clone #47929) using the mMessage mMachine SP6 Transcription Kit. The RNA was purified with the QIAGEN RNeasy Mini Kit, and its concentration was measured using a Nanodrop spectrophotometer, followed by verification on a 3% agarose gel.

For zebrafish injections, fertilized one-cell stage embryos (WT-TL strain) were injected with approximately 800 ng of nCas9 mRNA, 400 ng of gRNA, and 1 µM of STOP cassette oligonucleotides. This procedure resulted in a zebrafish strain lacking functional TMEM16B protein, which was confirmed by the absence of I_Cl[Ca]_ currents, as measured in electrophysiological recordings (Figure 1).

### Electrophysiological recordings

#### Retina isolation

On the day of the experiment, zebrafish were dark-adapted for 30 minutes in a completely light-tight chamber. They were then euthanized by immersion in iced water, decapitated, and their eyes were enucleated. Retinas were isolated under dim red illumination and positioned with the photoreceptor side facing up in a recording chamber (300 μl; model RC-26G, Warner Instruments) mounted on a Nikon Eclipse 600FN microscope. The preparation was observed on an LCD monitor using a CCD camera and infrared illumination (λ > 800 nm), with a ×60 water-immersion objective (NA 1.0) and differential interference contrast optics (Kodak Wratten filter 87c, USA).

#### Recording solutions

##### Extracellular solution

The tissue was continuously superfused with oxygenated Ringer’s solution at room temperature (20°C). The solution had the following composition (in mM): 102.0 NaCl, 2.6 KCl, 1.0 MgCl₂, 1.0 CaCl₂, 28.0 NaHCO₃, and 5.0 glucose. It was constantly gassed with a mixture of 2.5% CO₂ and 97.5% O₂ to maintain a pH of 7.8, with an osmolarity of 245–255 mosmol l⁻¹.

##### Intrapipette solution

The patch pipette solution contained (mM): 96 potassium gluconate, 10 KCl, 1 MgCl_2_, 0.036 CaCl_2_ ,0.075 EGTA, 5 HEPES, 5 ATP-K_2_,1 GTP-Na_3_, 0.1 cGMP-Na, 20 phosphocreatine-Na_2_ and 50 units ml−1 creatine phosphokinase, adjusted with KOH to pH 7.27–7.3 (osmolarity 265–275 mosmol l^−1^). The EGTA concentration was chosen to match estimates of the endogeneous calcium buffering in the photoreceptors, that was found to be analogous to 0.05-0.1mM (35,36). The CaCl_2_ concentration was chosen to ensure that the resulting concentration of free Ca^2+^ matched endogenous [Ca^2+^] in photoreceptors in the dark (∼300nM (48)). E_cl_ was calculated −55mV, in accordance with measurements performed in ref. (18).

All chemicals were supplied by Sigma-Aldrich (Zwijndrecht, The Netherlands), except for NaCl (Merck Millipore, Amsterdam, The Netherlands).

#### Recording procedure

Measurements from zebrafish cones were conducted in current-clamp mode (for voltage light responses and dark potential resistance measurements) and voltage-clamp mode (for I_Cl[Ca]_ and its reversal potential). Patch pipettes (resistance: 8–12 MΩ, PG-150T-10; Harvard Apparatus, Holliston, MA, USA) were pulled using a Brown Flaming Puller (model P-87; Sutter Instruments, Novato, CA, USA).

Filled patch pipettes were mounted on an MP-85 Huxley/Wall-type manual micromanipulator (Sutter Instrument) and connected to a HEKA EPC-10 Dual Patch Clamp amplifier (HEKA Elektronik, Lambrecht, Germany). After achieving the whole-cell configuration, cones were spectrally classified based on their response amplitudes to long-, mid-, or short-peak wavelength light flashes (see Stimuli subsection). Subsequent light stimuli were delivered using only the light source that elicited the strongest response during spectral classification. Only cells exhibiting stable and rapid light responses were used for recording light-induced voltage responses. However, cells that did not respond to light were still included for measurements of I_Cl[Ca]_, its reversal potential, and cone dark potential resistance.

Data were acquired at a sampling rate of 200 Hz for light responses and 1 kHz for electrophysiological properties using the Patchmaster software package (HEKA Elektronik).

### OKR setup

Zebrafish larvae (7 dpf) were carefully transferred from the petri dish using a disposable pipette and placed in a 15 mm petri dish containing 3% methylcellulose. To maintain the methylcellulose’s ability to restrict involuntary movements, a minimum of E3 medium was carried over during the transfer. Each larva was positioned at the center of the dish with its dorsal side facing up, ensuring both eyes were aligned at the same level. If necessary, gentle adjustments were made to achieve the correct orientation.

The petri dish containing the zebrafish larva was then placed into the experimental setup (Figure 6). The setup, previously described in detail by Klaassen et al. (59), consists of an LED-based optokinetic stimulator equipped with a built-in camera and two PCs. The stimulator comprises 60 circularly arranged plexiglass slides, each illuminated by 10 LEDs of three different wavelengths (Figure 6). The LED intensity can be adjusted every 4 milliseconds, allowing the generation of a moving grating stimulus with sinusoidal properties at varying velocities and contrasts. Infrared (IR) light illuminates the fish, enabling eye movement tracking via an IR camera.

### Stimuli

#### Light Stimuli

The light stimulator was a custom-built LED system incorporating a three-wavelength high-intensity LED (Atlas, Lamina Ceramics, Westhampton, NJ, USA). The LEDs had peak emissions at 465, 525, and 624 nm, with a bandwidth of less than 25 nm. Linearity was maintained via an optical feedback loop. The LED output was delivered to the microscope through an optical fiber and focused onto cone outer segments using a ×60 water-immersion objective. The mean light intensity across all stimuli ranged from 0.12 to 1.84 × 10⁵ photons μm⁻² s⁻¹, varying according to the amplitude of light steps at 465, 525, and 624 nm, which were used for cone spectral classification. We measured cone voltage responses using two stimulus types: Weber contrast steps (Figure 2) and a sum of sinewaves (Figure 3). The stimuli were presented at the frequency of 200Hz.

##### Weber contrast steps

The stimulation protocol is visualized in Figure 2. Contrast steps had an amplitude of 0.8 Weber contrast units, calculated with equation 1:

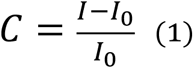

where C – is Weber contrast, I - is the light intensity during the contrast step, I_0_ - is the mean light intensity before the contrast step. The duration of the contrast steps was 500ms.

##### Sum of sinewaves

Sum of sinewaves stimuli (inspired by Shapley and Victor, 1978; Howlett et al., 2017), were 10 seconds long and consistent of the sum of 21 sinewaves with randomized phases. The sinewave frequencies were chosen to mimic the power spectra distribution of natural scenes, where most of the power lies in a lower frequency end of the spectrum (42,61,62). Such naturalistic stimuli were shown to be crucial for revealing non-linear properties of the cone photoreceptors (2,39). The employed frequencies were (in Hz): 0.2, 0.3, 0.5, 0.7, 1.1, 1.3, 1.9, 2.3, 3.1, 4.1, 5.3, 6.1, 7.3, 8.9, 10.7, 13.9, 16.7, 19.6, 23.3, 27.4, 31.7.

We used two types of the sum of sinewaves stimuli: high and low contrast. High contrast stimuli had r.m.s. contrast of 0.32, low contrast stimuli had r.m.s. contrast of 0.063.

Weber contrast steps and sum of sinewaves stimuli were recorded within one 23.5 seconds long protocol.

#### Electrical stimuli

To characterize electrophysiological properties of I_Cl[Ca]_ we performed four types of experiments: measurement of the tail current (Figure 1A, F), measurement of the tail current reversal potential (Figure 1C), measurement of the dark resting-membrane potential and measurement of the cone dark potential-resistance (Figure 1H). The electrical stimuli were delivered and measured at frequency of 1kHz.

To measure the tail current, we clamped the cone potential at a nominal (i.e. before series resistance R_S_ compensation) value of −70mV for 1500ms. Then we performed 50ms step to the potential of −65mV to estimate R_S_ between the electrode and the cell (63). After R_S_ measurements, we held the cell at −70mV for additional 450ms before depolarizing it for 2000ms to the nominal potential between −60 to 10 mV (in steps of 10mV). After this we stepped back to −70mV and recorded 2000ms of the tail current.

The stimulus to measure the reversal potential of the tail current was similar, with two exceptions. First, we always depolarized cone to the potential of 10mV. Secondly, to measure tail current we stepped the cone membrane potential to the potentials in the range from −60 to 10mV.

To measure the cone dark membrane potential, we performed one-minute-long current clamp recording with the current clamped at zero.

To measure cone dark potential resistance, we performed current clamp recordings where from the zero current baseline cones we injected a current step of 5pA for 50ms, followed by the 300ms holding at zero current, followed by the current step of −5pA for another 50ms.

#### OKR Stimuli

The optokinetic response (OKR) was measured in zebrafish larvae exposed to various moving grating stimulus with different angular velocities (degrees per second) and spatial frequencies (cycles per degree). All stimuli were presented with a Michelson contrast of 100%, calculated with equation 2:

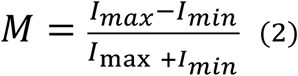

Where M is Michelson contrast, I_max_ is the maximal light intensity within the sequence (white bar of the grating), I_min_ is the minimal intensity within the sequence (dark bar of the grating).

The velocities tested were: 2, 5, 9, 12, 16, and 25 degrees per second. Each velocity was paired with different spatial frequencies: 0.014, 0.033, and 0.056 cycles per degree, resulting in a total of 18 unique stimulus combinations. To prevent habituation, the stimulus direction was periodically alternated. Eye movements of the zebrafish larvae were recorded using an infrared camera to assess the OKR.

### Data Analysis

#### Light responses

##### Preprocessing stage

In vitro recordings are recordings in a dying tissue. In the retina, this issue is further exacerbated because the retinal pigment epithelium, which resupplies photoreceptors with retinal, is removed during dissection. Therefore, when recording light responses, it is essential to ensure stable recording conditions.

To systematically exclude unstable recordings, we applied three primary criteria:

- The standard deviation of an individual response exceeded two standard deviations of the mean trace’s standard deviation within the same recording condition (i.e., recordings at the same light intensity in an individual cone). A two-standard-deviation range captures 95% of the data, a commonly used confidence interval (CI) in statistical tests (Colquhoun, n.d.).
- The trace contained noise spikes with amplitudes exceeding 20 mV, which could bias response amplitude estimates and indicate compromised cell health.
- The absence of an observable response in the high-contrast portion of the stimulus, suggesting insufficient photopigment levels for a light response.

Additionally, we calculated noise power (see below) in the recorded cones and excluded two photoreceptors (one WT, one mutant) whose total noise power amplitude were at least 10 times higher than in any other cone.

##### Photoreceptor signal transfer properties

To characterize the transfer properties of cone photoreceptors, we computed the transfer function from the recorded voltage responses to sum of sinewaves stimuli. The analysis was performed using custom Python scripts based on Fourier methods.

First, both stimulus (S(t)) and response (R(t)) signals were linearly detrended to remove the slow drifts. Then the cross-spectral density (P_SR_(f)) between the stimulus and response was estimated using Welch’s method (64) with a rectangular window of 2000 data points (the entire response stretch) and no overlap. The power spectral density of the stimulus (P_SS_(f)) was estimated in the same way. The transfer function (T(f)) was then obtained as the ratio of P_SR_(f)/P_SS_(f).

Note, that a rectangular window was appropriate in this case because the stimuli were the sum of properly spaced sinewaves. The total stimulus duration (10 seconds) ensured sufficient frequency resolution, and we only analyzed frequencies that were explicitly present in the stimulus. This minimized spectral leakage, making more complex windowing functions unnecessary.

To compute the impulse response function, we linearly interpolated transfer function between sum of sinewaves frequencies (to obtain smooth estimation) and performed an inverse Fourier transform of the transfer function. We extracted the real part of the inverse Fourier transform.

We verified the applicability of linear system analysis for the zebrafish cone photoreceptors by estimating correlation between recorded voltage responses and voltage response obtained by convolution of impulse response function with the stimuli. The obtained correlation values were in the range of 75%-99% and always higher than 90% for the high contrast stimuli. Thus, we concluded that linear system analysis was appropriate.

To compute Signal-to-Noise Ratio (SNR) we assumed that cone voltage responses averaged across individual traces are signals, whereas differences between individual traces and mean trace was considered to be noise. To estimate signal power spectrum, we used Welch’s approach as described above. With the same method we calculate power spectra of the individual noise traces and then averaged them together to obtain noise power spectrum for each of the cone photoreceptors. SNR was computed with equation 3 following the approach by Van Hateren and Snippe(65):

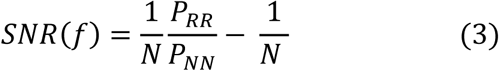

Where N is a number of recorded traces.

To estimate information transfer spectra and information channel capacity (IC) we employed approach developed by Shannon(66), equation 4:

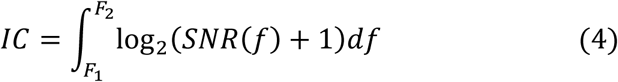

Where F_1_ and F_2_ are limits of the frequencies in the sum of sinewave stimuli.

##### I_Cl[Ca]_ electrophysiological properties

As was described in the section about recording procedures, voltage-clamp protocols to measure I_Cl[Ca]_ (Figure 1) contained 50ms step with amplitude of 5mV used to determine series resistance R_S._ This was done to subsequently adjust nominal voltage steps values for a R_S_ to obtain true voltage values that affected cone photoreceptors. The R_S_ was calculated as the ratio between depolarization amplitude and transient current response.

To estimate reversal potential of the I_Cl[Ca]_ (Figure 1C) we start with calculation of the integral under the tail current to obtain tail current charge transfer at different holding potentials. Then we fitted linear regression between the transferred charge and holding potential to find the potential where charge is zero i.e. no current was flowing through the corresponding channel. This potential was the reversal potential.

For the measurements of dark potential resistance, we estimated the amplitude of the voltage response upon ±5pA current injection as mean potential averaged through the last 20ms of the current step. Since there were two current steps, we estimated cone resistance as mean between corresponding absolute changes in the membrane potential.

### Statistics

Before performing any statistical tests, we excluded outliers using boxplots. According to the common convention, we considered a data point as an outlier if its corresponding value was 1.5*IQR (Inter-Quantile Range) larger than the 75^th^ (Q3) percentile or 1.5*IQR smaller than the 25^th^ (Q1) percentile of dataset. IQR – the range between Q3/Q1 and dataset’s median.

To test for significance, we employed two-sample t-test (except for Figure 2) The data is presented as mean ± SEM (Standard Error of the Mean), unless otherwise specified.

For the Figure 2 (OKR data), since we had repeated measurements from the same fish (i.e., multiple velocities and spatial frequencies from each animal), we fitted the data with a linear mixed-effects model to account for the non-independence of repeated measures and the influence of both stimulus velocity and spatial frequency on the OKR gain. The model included both a random intercept and a random slope for spatial frequency to better capture within-subject variability.

The analysis included *N* = 43 observations obtained from 17 individual fish, with fish identity modelled as a random effect. Fixed effects included genotype (wild-type vs mutant) and spatial frequency (three levels). The model was fitted in Python using the statsmodels package with restricted maximum likelihood estimation and the LBFGS optimizer.

## Notes

### Competing Interest Statement

The authors have declared no competing interest.

### Summary of Updates

This version of the manuscript was revised to show the parallels between the calcium-dependent chloride current function in vision and odorant system

### Funding

- European Union: Horizon 2020: 674901

## References

1. Kamar S, Howlett MHC, Kamermans M. Silent-substitution stimuli silence the light responses of cones but not their output. J Vis. 2019;19(5):1–11.

2. Howlett MHC, Smith RG, Kamermans M. A novel mechanism of cone photoreceptor adaptation. PLoS Biol [Internet]. 2017;15(4):e2001210. Available from: http://www.ncbi.nlm.nih.gov/pubmed/28403143%0Ahttp://dx.doi.org/10.1371/journal.pbio.2001210

3. Wen X, Thoreson WB. Contributions of glutamate transporters and Ca2+-activated Cl− currents to feedback from horizontal cells to cone photoreceptors. Exp Eye Res [Internet]. 2019;189(August):107847. Available from: 10.1016/j.exer.2019.107847

4. Vroman R, Kamermans M. Feedback-induced glutamate spillover enhances negative feedback from horizontal cells to cones. J Physiol. 2015;593:2927–2940.

5. Lalonde MR, Kelly ME, Barnes S. Calcium-activated chloride channels in the retina. Channels. 2008;2(4):252–60.

6. van Hateren H. A cellular and molecular model of response kinetics and adaptation in primate cones and horizontal cells. J Vis. 2005;5(4):331–47.

7. Fahey PK, Burkhardt DA. Effects of light adaptation on contrast processing in bipolar cells in the retina. Vis Neurosci. 2001;18(4):581–97.

8. Thoreson WB, Burkhardt DA. Ionic influences on the prolonged depolarization of turtle cones in situ. J Neurophysiol. 1991;65(1):96–110.

9. Steven Barnes; Hille Bertil. Ionic channels of the inner segment of tiger salamander cone photoreceptors. J Gen Physiol. 1989;94(4):719–43.

10. Barnes S. After transduction: Response shaping and control of transmission by ion channels of the photoreceptor inner segment. Neuroscience. 1994;58(3):447–59.

11. Barnes S. Ionic channels of the inner segment of tiger salamander cone photoreceptors. J Gen Physiol. 2004;94(4):719–43.

12. Baylor DA, Fuortes MGF, O’Bryan PM. Receptive fields of cones in the retina of the turtle. J Physiol. 1971;214(2):265–94.

13. O’Bryan PM. Properties of and the Mechanisms Underlying the Horizontal-Cell-Mediated Depolarization of Cones Which Is Evoked By Surround Illumination. 1973;207–23.

14. Burkhardt DA, Gottesman J, Thoreson WB. Prolonged depolarization in turtle cones evoked by current injection and stimulation of the receptive field surround. J Physiol [Internet]. 1988 Dec 1;407(1):329–48. Available from: 10.1113/jphysiol.1988.sp017418

15. Maricq AV, Korenbrot JI. Calcium and calcium-dependent chloride currents generate action potentials in solitary cone photoreceptors. Neuron. 1988;1(6):503–15.

16. Lasansky A. Synaptic action mediating cone responses to annular illumination in the retina of the larval tiger salamander. J Physiol [Internet]. 1981 Jan 1;310(1):205–14. Available from: 10.1113/jphysiol.1981.sp013544

17. Barnes S, Bui Q. Modulation of calcium-activated chloride current via pH-induced changes of calcium channel properties in cone photoreceptors. Journal of Neuroscience. 1991;11(12):4015–23.

18. Kraaij DA, Spekreijse H, Kamermans M. The nature of surround-induced depolarizing responses in goldfish cones. Journal of General Physiology. 2000;115(1):3–15.

19. Verweij J, Hornstein EP, Schnapf JL. Surround Antagonism in Macaque Cone Photoreceptors. Journal of Neuroscience. 2003;23(32):10249–57.

20. Yagi T, Macleish PR. Ionic conductances of monkey solitary cone inner segments. J Neurophysiol. 1994;71(2):656–65.

21. Frederiksen R, Bonezzi PJ, Fain GL, Sampath AP. The Role of the Ca2+-activated Cl− Conductance in the Membrane Potential and Light Response of Mouse Rods. Journal of Neuroscience. 2025;45(22):1–11.

22. Koch C. Biophysics of Computation: Information Processing in Single Neurons [Internet]. Oxford University Press; 1998. Available from: 10.1093/oso/9780195104912.001.0001

23. Hille B. Ion Channels of Excitable Membranes. Vol. 18, Ion Channels of Excitable Membranes. 2001. 1–814 p.

24. Zang J, Neuhauss SCF. The binding properties and physiological functions of recoverin. Front Mol Neurosci. 2018;11(December):1–10.

25. Vroman R, Klaassen LJ, Howlett MHC, Cenedese V, Klooster J, Sjoerdsma T, et al. Extracellular ATP Hydrolysis Inhibits Synaptic Transmission by Increasing pH Buffering in the Synaptic Cleft. PLoS Biol [Internet]. 2014 May 20;12(5):e1001864. Available from: 10.1371/journal.pbio.1001864

26. Dibattista M, Pifferi S, Hernandez-Clavijo A, Menini A. The physiological roles of anoctamin2/TMEM16B and anoctamin1/TMEM16A in chemical senses. Cell Calcium [Internet]. 2024;120(February):102889. Available from: 10.1016/j.ceca.2024.102889

27. Reisert J, Pifferi S, Guarneri G, Ricci C, Menini A, Dibattista M. The Ca2+-activated Cl− channel TMEM16B shapes the response time course of olfactory sensory neurons. Journal of Physiology. 2024;602(19):4889–905.

28. Stöhr H, Heisig JB, Weber BHF, Schulz HL, Schöberl S, Milenkovic VM, et al. TMEM16B, a novel protein with calcium-dependent chloride channel activity, associates with a presynaptic protein complex in photoreceptor terminals. Journal of Neuroscience. 2009;29(21):6809–11.

29. Mercer AJ, Rabl K, Riccardi GE, Brecha NC, Stella SL, Thoreson WB. Location of release sites and calcium-activated chloride channels relative to calcium channels at the photoreceptor ribbon synapse. J Neurophysiol. 2011;105(1):321–35.

30. Endeman D, Fahrenfort I, Sjoerdsma T, Steijaert M, ten Eikelder H, Kamermans M. Chloride currents in cones modify feedback from horizontal cells to cones in goldfish retina. Journal of Physiology. 2012;590(22):5581–95.

31. Scudieri P, Sondo E, Ferrera L, Galietta LJV. The anoctamin family: TMEM16A and TMEM16B as calcium-activated chloride channels. Exp Physiol. 2012;97(2):177–83.

32. Jeon JH, Paik SS, Chun MH, Oh U, Kim IB. Presynaptic Localization and Possible Function of Calcium-Activated Chloride Channel Anoctamin 1 in the Mammalian Retina. PLoS One. 2013;8(6).

33. Ponissery Saidu S, Stephan AB, Talaga AK, Zhao H, Reisert J. Channel properties of the splicing isoforms of the olfactory calcium-activated chloride channel Anoctamin 2. Journal of General Physiology. 2013;141(6):691–703.

34. Betto G, Cherian OL, Pifferi S, Cenedese V, Boccaccio A, Menini A. Interactions between permeation and gating in the TMEM16B/anoctamin2 calcium-activated chloride channel. J Gen Physiol. 2014;143(6):703–18.

35. van Hook MJ, Thoreson WB. Weak endogenous Ca2+ buffering supports sustained synaptic transmission by distinct mechanisms in rod and cone photoreceptors in salamander retina. Physiol Rep. 2015;3(9):1–22.

36. Van Hook MJ, Thoreson WB. Endogenous calcium buffering at photoreceptor synaptic terminals in salamander retina. Synapse. 2014;68(11):518–28.

37. Pedemonte N, Galietta LJ V. Structure and Function of TMEM16 Proteins (Anoctamins). Physiol Rev. 2014;94(2):419–59.

38. Thoreson WB, Bryson EJ. Chloride equilibrium potential in salamander cones. BMC Neurosci. 2004;5:1–7.

39. Yedutenko M, Howlett MHC, Kamermans M. Enhancing the dark side: asymmetric gain of cone photoreceptors underpins their discrimination of visual scenes based on skewness. Journal of Physiology. 2022;600(1):123–42.

40. Baccus SA, Meister M. Fast and slow contrast adaptation in retinal circuitry. Neuron. 2002;36(5):909–19.

41. Schreyer HM, Gollisch T. Nonlinear spatial integration in retinal bipolar cells shapes the encoding of artificial and natural stimuli. Neuron [Internet]. 2021;109(10):1692–1706.e8. Available from: 10.1016/j.neuron.2021.03.015

42. Van Hateren JH. Processing of natural time series of intensities by the visual system of the blowfly. Vision Res. 1997;37(23):3407–16.

43. Guarneri G, Pifferi S, Dibattista M, Reisert J, Menini A, Area N, et al. Paradoxical electro-olfactogram responses in TMEM16B knock-out mice. 2023;(January):1–8.

44. Rinner O, Rick JM, Neuhauss SCF. Contrast sensitivity, spatial and temporal tuning of the larval zebrafish optokinetic response. Invest Ophthalmol Vis Sci. 2005;46(1):137–42.

45. Wald A. Tests of Statistical Hypotheses Concerning Several Parameters When the Number of Observations is Large. Trans Am Math Soc. 1943;54(3):426–82.

46. Schoukens J, Pintelon R, Der Ouderaa E Van, Renneboog J. Survey of Excitation Signals for FFT Based Signal Analyzers. IEEE Trans Instrum Meas. 1988;37(3):342–52.

47. Pifferi S, Dibattista M, Menini A. TMEM16B induces chloride currents activated by calcium in mammalian cells. Pflugers Arch. 2009;458(6):1023–38.

48. Jackman SL, Choi SY, Thoreson WB, Rabl K, Bartoletti TM, Kramer RH. Role of the synaptic ribbon in transmitting the cone light response. Nat Neurosci. 2009;12(3):303–10.

49. Okada T, Horiguchi H, Tachibana M. Ca2+-dependent C− current at the presynaptic terminals of goldfish retinal bipolar cells. Neurosci Res [Internet]. 1995;23(3):297–303. Available from: https://www.sciencedirect.com/science/article/pii/0168010295009558

50. Zhang Y, Zhang Z, Xiao S, Tien J, Le S, Le T, et al. Inferior Olivary TMEM16B Mediates Cerebellar Motor Learning. Neuron [Internet]. 2017;95(5):1103–1111.e4. Available from: 10.1016/j.neuron.2017.08.010

51. Sterling P, Laughlin S. Principles of Neural Design [Internet]. The MIT Press; 2015. Available from: http://www.jstor.org/stable/j.ctt17kk982

52. Yedutenko M, Howlett MHC, Kamermans M. High Contrast Allows the Retina to Compute More Than Just Contrast. Front Cell Neurosci. 2021;14(January):1–19.

53. Korenbrot JI, Rebrik TI. Tuning outer segment Ca2+ homeostasis to phototransduction in rods and cones. Adv Exp Med Biol. 2002;514:179–203.

54. Arshavsky VY, Burns ME. Photoreceptor signaling: Supporting vision across a wide range of light intensities. Journal of Biological Chemistry [Internet]. 2012;287(3):1620–6. Available from: 10.1074/jbc.R111.305243

55. Fain GL, Matthews HR, Cornwall MC, Koutalos Y. Adaptation in vertebrate photoreceptors. Physiol Rev. 2001;81(1):117–51.

56. Choi SY, Borghuis B, Rea R, Levitan ES, Sterling P, Kramer RH. Encoding light intensity by the cone photoreceptor synapse. Neuron. 2005;48(4):555–62.

57. Borghuis BG, Ratliff CP, Smith RG. Impact of light-adaptive mechanisms on mammalian retinal visual encoding at high light levels. J Neurophysiol. 2018;119(4):1437–49.

58. Hwang WY, Fu Y, Reyon D, Maeder ML, Tsai SQ, Sander JD, et al. Efficient genome editing in zebrafish using a CRISPR-Cas system. Nat Biotechnol [Internet]. 2013;31(3):227–9. Available from: 10.1038/nbt.2501

59. Klaassen LJ, Sun Z, Steijaert MN, Bolte P, Fahrenfort I, Sjoerdsma T, et al. Synaptic transmission from horizontal cells to cones is impaired by loss of connexin hemichannels. PLoS Biol. 2011;9(7).

60. Shapley RM, Victor JD. The effect of contrast on the transfer properties of cat retinal ganglion cells. J Physiol. 1978;(285):275–98.

61. Van Hateren JH, Rüttiger L, Sun H, Lee BB. Processing of natural temporal stimuli by macaque retinal ganglion cells. Journal of Neuroscience. 2002;22(22):9945–60.

62. Balboa RM, Grzywacz NM. Power spectra and distribution of contrasts of natural images from different habitats. Vision Res. 2003;43(24):2527–37.

63. Ogden D, Stanfield P. Patch clamp techniques for single channel and whole-cell recording. Currents [Internet]. 1981;2(7):53–78. Available from: http://www.utdallas.edu/∼tres/microelectrode/microelectrodes_ch04.pdf

64. Welch PD. Welchs Periodogram.pdf. Vol. 15, IEEE Trans. Audio and electroacoustic. 1967. p. 70–3.

65. van Hateren JH, Snippe HP. Information theoretical evaluation of parametric models of gain control in blowfly photoreceptor cells. Vision Res [Internet]. 2001;41(14):1851–65. Available from: pm:11369048%5Cnhttp://www.sciencedirect.com/science/article/pii/S0042698901000529

66. Shannon CE. A Mathematical Theory of Communication. 1948;27(April 1928):379–423.

